# Dynamic longitudinal behavior in animals exposed to chronic social defeat stress

**DOI:** 10.1101/2020.01.17.907477

**Authors:** M. Wendelmuth, M. Willam, H. Todorov, K. Radyushkin, S. Gerber, S. Schweiger

**Affiliations:** Institute for Human Genetics, University Medical Center, Mainz; Leibniz Institute for Resilience Research, Mainz; Computational Systems Genetics, Faculty of Biology, Institute for Developmental Biology and Neurobiology (IDN) and Center for Computational Sciences in Mainz, Johannes Gutenberg-University Mainz

## Abstract

Chronic social defeat (CSD) can lead to impairments in social interaction and other behaviors that are supposed to model features of major depressive disorder (MDD). Not all animals subjected to CSD, however, develop these impairments, and maintained social interaction in some animals is widely used as a model for resilience to stress-induced mental dysfunctions. So far, animals have mainly been studied shortly (24 hours and 7 days) after CSD exposure and longitudinal development of behavioral phenotypes in individual animals has been mostly neglected. We have analyzed social interaction and novel object recognition behavior of stressed mice at different time points after CSD and have found very dynamic courses of behavior of individual animals. Instead of the two groups, resilient or susceptible, that are found at early time points our data suggest four groups with (i, ii) animals behaving resilient or susceptible at early and late time points, respectively (iii) animals that start susceptible and recover with time or (iv) animals that are resilient at early time points but develop vulnerability later on.

## Introduction

Major depressive disorder (MDD) is one of the leading causes of disability in Western societies (1). While chronic stress exposure does significantly contribute to its pathogenesis, not everybody exposed to chronic stress develops MDD (2).

It has been suggested that changes in animal behavior induced by chronic social defeat (CSD) are suitable models for MDD (3–5). In this model, animals are exposed to an aggressor mouse every day for a couple of minutes (5-10 min) over a timespan of 10-21 days. During the rest of the day they are housed in the same cage as the aggressor but separated by a transparent wall (suppl. Fig. 1). Subsequent behavior analysis revealed avoidance behavior in social interaction tests in many of the tested animals. Several animals however did not show aversive reaction 1-7 days after CSD exposure, suggesting resilience.

Resilience has often been described as a static personality trait that protects from aversive stress reactions - a yes or no in the ability of individuals to defend themselves. However, it becomes more and more clear that resilience is the result of a dynamic process over time that leads to successful adaptation to stressors. This has recently been summarized in a concept paper by Kalisch et al (6), who have drafted a “Hybrid Symptom and Resilience Factor model” describing psychiatric symptoms in a well-connected network influencing each other and resilience factors as nodes that at variable anchor points interfere with the network. This however means that at different timepoints the networks may arrive at individually different stages, swapping either into a vulnerable, susceptible status or reaching a point of adaptation and functionality.

Using the CSD model we hypothesized that, rather than having a fixed phenotype after a defined timepoint with animals either belonging into the susceptible or the resilient group, a dynamic process of changes between affected behavior and recovery on the way to adaptation will be visible. So far, mice exposed to CSD have extensively been used to describe and analyze susceptible and resilient behavior at a static time point after the last defeat. Longitudinal development of behavior after CSD in individual animals however still mostly remains unexplored in animals. In this study, we monitored different domains of aberrant behavior (social interaction and novel object recognition) after CSD in different groups of animals tested at 24 hours, 7 days or 21 days after CSD. We then went on testing one and the same group of animals after CSD 7 days, 21 days and 42 days after the last defeat and followed individual animals in their behavior process.

## Materials and methods

### Animals

Male C57BL/6J mice arrived at 11 weeks of age in our animal facility. The mice were group-housed in standard cages (2–4 mice per cage) and acclimatized to a temperature- and humidity-controlled room (temperature = 22 ± 2 °C, relative humidity = 50 ± 5%) for at least 7 days prior to being moved to a new cage and presented with an unknown CD1 mouse to induce social defeat (see below). The mice were maintained under a 12-h light/dark cycle with *ad libitum* access to food and water. The groups consisted of C57BL/6J mice weighing 25–30g without any apparent injuries or loss of voluntary movements. C57BL/6J mice were assigned to two groups on the first day of social defeat (Day-10). This study was conducted in accordance with the European directive 2010/63/EU for animal experiments and was approved by the local authorities (approval number G17-1-049, Animal Protection Committee of the State Government, Landesuntersuchungsamt Rheinland-Pfalz, Koblenz, Germany). Retired male CD1 breeder mice aged 6–12 months were used as aggressors and housed individually until screening. Screening for aggressiveness was performed on the day before the first day of defeat. All C57BL/6J mice used in this study were purchased from Janvier (France).

### Social defeat

A CSD stress protocol was carried out using the resident/intruder paradigm described in previous reports (3, 7, 8), with some modifications. In short, every experimental mouse was placed in a home cage of a CD1 (aggressor) mouse and experienced attack bites (contact phase). After the defeat episode, the two mice were separated by a metal mesh and housed in the same cage for 24 h to expose the subordinate mouse to sensory stress without having physical contact. The defeat cycle was repeated for 10 consecutive days. To prevent habituation the resident/intruder pair was changed every day (suppl. Fig. 1). To limit stress and physical injury to the intruder mice to an acceptable level, we modified the protocol by restricting the duration of physical contact to 15-25s. As a control, non-defeated (non-stressed) mice were pair-housed in divided cages with no physical contact between the mice. After the last day of defeat all experimental and control mice were single-housed.

### Social interaction test

Social interaction (SI) testing was carried out as it was described in previous reports (1, 5, 6) with slight modifications. Each mouse was placed in an open field box (40 cm wide × 40 cm deep × 40 cm high), that had a cylindrical wire-mesh enclosure at 1 side. Dim lighting (50 lux) was used and the mouse was allowed to move freely in the field. Each test consisted of 2 sessions abiding 150 s. In the first session the wire-mesh enclosure was empty. After a short break of 30 s the second session started, in which an aggressive CD1 mouse that was novel to the C57BL/6J was placed within the wire-mesh enclosure (suppl. Fig. 2).

The exploratory behaviors of the mice were recorded with a charge-coupled device (CCD) camera and analyzed with video tracking software EthoVision XT (Noldus Information Technology). The amount of time each mouse spent in a round area (8 cm radial from mesh) surrounding the enclosure, defined as the interaction zone (IZ), was determined and the SI index was calculated as: (time mouse spent in the IZ when the target was present) / (time mouse spent in the IZ when target was absent) multiplied with 100. Scores < 100 were considered to be indicators of social avoidance behavior. After the second session, the field was cleaned with wet paper towels and the test session was repeated with a different mouse.

### 24 hours Novel object recognition test

For the first sample phase of the Novel object recognition test (NORT), mice were placed into an arena (40 cm wide × 40 cm deep × 40 cm high) and presented with two identical objects. Mice were allowed to explore the arena and the objects for 5 minutes before being removed for a 24 h interval spent in their home cage. Mice were then given a second 5 min sample phase. Following the second sample phase, the mice were returned to their home cage for another 24h retention interval. In the test phase on the third day, one of the items was replaced with a novel object (suppl. Fig. 3). In the test phase behavior of mice was recorded with a charge-coupled device (CCD) camera. The test phase continued for 10 minutes. The location of the objects and the object that was replaced with the novel item was fully counterbalanced both within and between groups. The videos were analyzed with BORIS v. 6.3.9 (9) and time for exploring both objects, familiar and unfamiliar, was determined. The discrimination Index (DI) was calculated as (time unfamiliar object) – (time familiar object) / (time unfamiliar object + time familiar object).

### Statistical analyses

Outliers were identified using the ROUT function (Q=1%) in GraphPad Prism. Normality of groups was checked by using D’Agostino & Pearson tests. All 2-group comparisons were calculated using either t-test, when data followed a Gaussian distribution, or Mann-Whitney test, when data were not normally distributed. For comparisons of three groups either ANOVA or Kruskal-Wallis test was performed accordingly. Post-hoc analysis was done using Dunn’s multiple comparisons test or Bonferroni’s multiple comparisons test. Correlations were analyzed by determining Pearson r. A 2-way analysis of covariance or a linear mixed effects model were used to investigate the effect of exploration behavior on DI scores. Statistical significance was defined as *p* < 0.05. Statistical analysis was performed with GraphPad Prism version 7.05 and the statistical language R version 3.5.3 (R Core Team, Vienna, Austria).

## Results

### Behavior changes over time after CSD

To investigate the consequences of CSD on social behavior and cognition over time, groups of test mice were exposed to CSD and analyzed in an SI test one day (T1), 7 days (T2) or 21 days (T3) after the last defeat. For cognition testing, mice were exposed to a NORT as described above also at T1-T3 (Fig. 1a). In order to avoid re-testing errors different mouse groups were used for the different time points.

While group behavior in the NORT at T1 did not show cognitive aberrations (Fig. 1e), the SI test (T1) revealed a significant decrease in social interaction with the aggressor for the stressed group (Fig. 1b), supporting previous findings (3, 7, 10–12). At T2 both SI and NORT exhibited significant deficits in the defeated group, indicated by less time spent in the interaction zone and decreased time spent with the novel object, respectively, compared to the control group (Fig 1c, f). Similar exploration times in all groups revealed that impairment in the NORTs is not caused by a decline in exploration motivation as has been suggested previously (T1-T3, suppl. Fig. 4a-c (13)).

**Figure 1:**
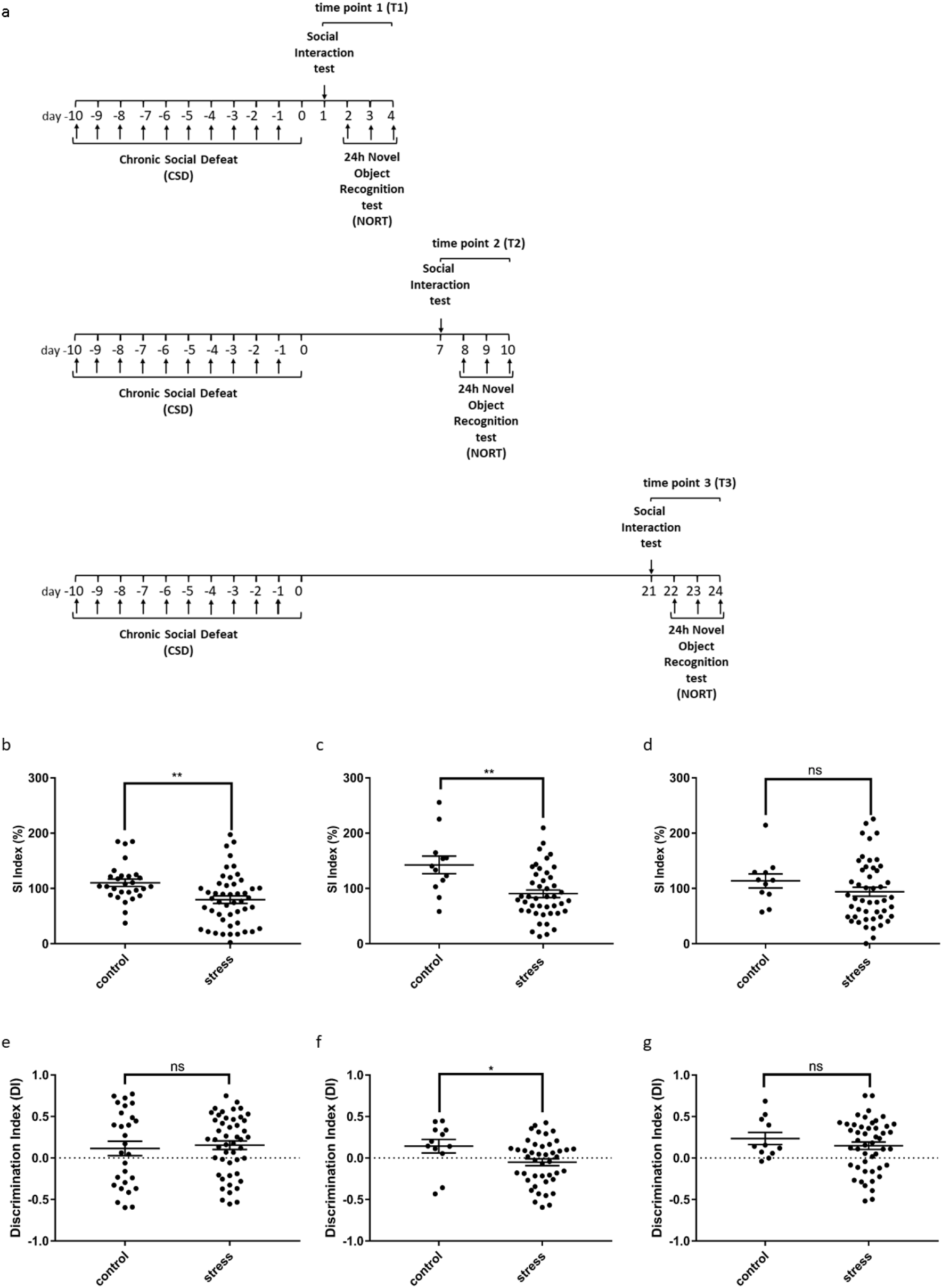
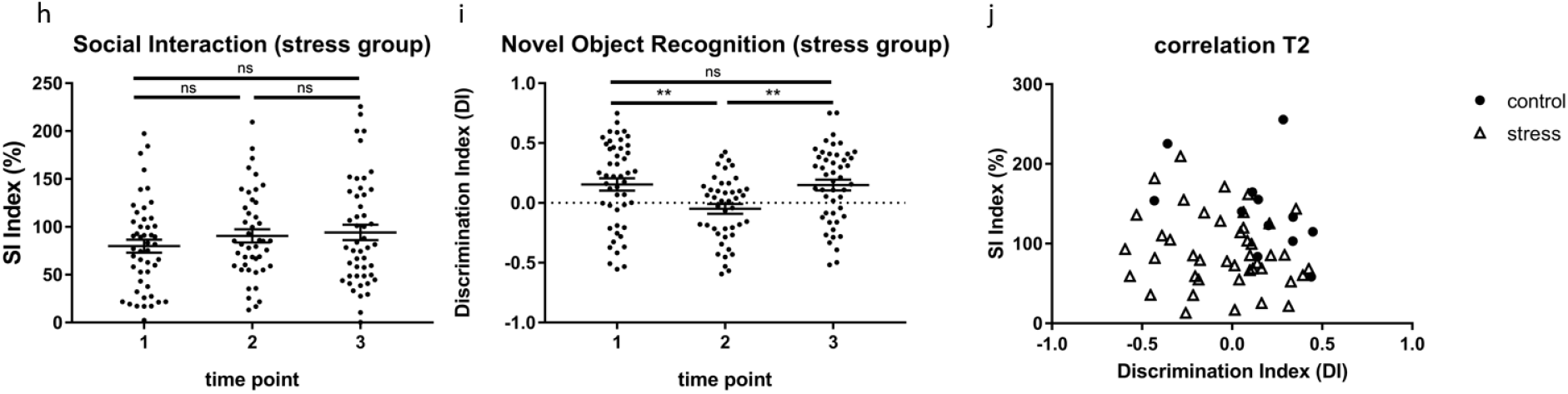
Animal behavior changes with time post exposure to CSD stress. (a) Experimental setup timeline. Within 11 days test mice were housed in sensory contact with a daily changed aggressor mouse. Each day they were exposed to fight for 15-25 seconds. Control mice were not exposed to fight, but housed equally. Each group was tested for social interaction and novel object recognition either 1-4 days (T1), 7-10 days (T2) or 21-24 days (T3) post CSD. (b-d) SI scores at T1, T2 or T3 of stressed and control mice. Social avoidance is clearly visible at T1 (p=0.0020, Unpaired t test with Welch’s correction) and T2 (p=0.0089, Unpaired t test with Welch’s correction). At T3 there is no significant difference between stressed and control mice, but still a tendency of social avoidance (p=0.2142, Unpaired t test with Welch’s correction). (e-g) DI scores after NORT of stressed and control mice at T1, T2 and T3. Shortly after CSD (T1) there was no significant difference between stressed and control mice (p=0.6979, Unpaired t test with Welch’s correction). At T2 stressed mice performed worse than control mice (p=0.0495, Unpaired t test with Welch’s correction). At T3 these deficits were abolished (p=0.4828, Mann-Whitney test). (h, i) Direct comparison of SI and DI scores from stressed mice of all three timepoints. (h) SI scores of stressed mice in direct comparison revealed no significant differences, suggesting a steady level of social avoidance (T1 vs.2 p=0.9217, T1 vs. 3 p=0.4893, T2 vs. T3 p=>0.9999, 1-way ANOVA with Bonferroni post hoc test) (i) DI scores of stressed mice in direct comparison demonstrated a significant decrease in cognitive skills at T2 with a full recovery at T3. (T1 vs.2 p=0.0049, T1 vs. 3 p=>0.9999, T2 vs. T3 p=0.0083, Kruskal-Wallis test and Dunn’s post hoc test) (j) Correlation of SI and DI scores at T2 reveals no significant association for individual mice (control r=−0.4457 p=0.1464, stress r=−0.1546 p=0.3163, Pearson r). Data are shown as mean ± standard error of the mean and individual values.

Surprisingly, at T3 the SI revealed slight social avoidance of the stressed group to the interaction zone compared to the control group, but without showing any significant difference between groups (Fig 1d). However, the control group tended to perform weaker than in the previous results, suggesting motivational bias in this group. By comparing social interaction performance of the stressed groups at all 3 time points (T1-T3), no obvious differences became visible, suggesting similarly impaired social interaction behavior of the T3 groups compared to the two other groups (Fig. 1h).

In contrast, the NORT at T3 revealed no significant difference between stressed and control group (Fig. 1g), indicating recovery of the stress-induced phenotype between T2 and T3. In direct comparison, DI scores of the NORT in the stressed group were significantly lower at T2 when compared with T1 and T3 (Fig. 1i). Hence, at T2 mice displayed deficits in cognitive abilities while the group at 21 days post CSD seemed to have recovered. No differences in exploration behavior were seen at any of the time points (suppl. Fig. 4). Furthermore, exploration time was not a significant predictor of DI scores in a 2-way analysis of covariance with experimental group and time as factors and exploration time as covariate (not shown). This confirmed that cognitive deficits at T2 in the stressed group were not due to reduced exploratory behavior.

Taken together, these results suggest a longitudinal lasting decrease in social interaction after CSD stressed compared to non-stressed mice. Also, novel object recognition seems to be affected, suggesting an involvement of cognitive domains after CSD exposure in stressed mice. However, this effect evolved in a delayed manner and did not last until T3 (quickly reversable).

Next, we asked, if impairment of SI correlates with object recognition decay at the T2 time point. We did not observe a significant overall correlation between the two behavioral domains for all animals pooled together. Furthermore, calculating Pearson r for control and stressed animals separately did not reveal a significant association between SI and DI scores (Fig. 1j). These results suggest independent development of social interaction and novel object recognition restriction after CSD exposure.

### Longitudinal evolvement of social interaction behavior changes in individual animals

We have shown how SI impairment develops 1 day (T1), 7 days (T2) and 21 days (T3) after exposure to CSD in different groups of animals. We then asked how SI behavior develops over time in individual animals. Thus, we exposed a group of 20 animals to CSD and then performed multiple testing of SI and NORT after 7 days (T2), 21 days (T3) and 42 days (T4) with the same animals (Fig. 2a). This repeated measures longitudinal design allowed us to not only track behavioral trajectories of single animals over time but also to statistically model inter-individual variability and exclude potential batch effects.

**Figure 2:**
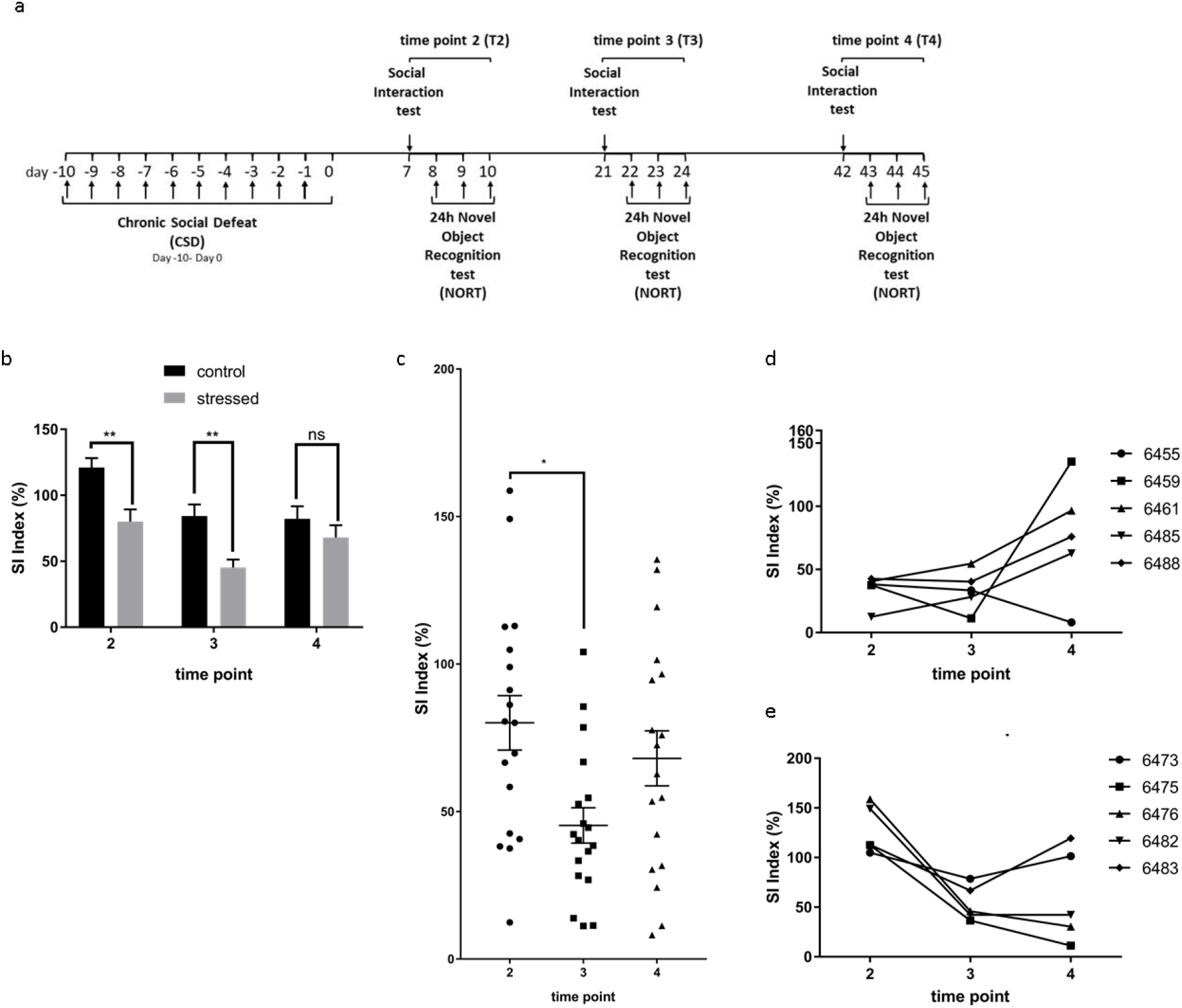
Longitudinal development of social interaction behavior after CSD stress. (a) Experimental setup timeline for retesting. Animals of a cohort of mice were (re-)tested for SI and NORT at 7-10 days (T2), 21-24 days (T3) and 42-45 days (T4) post CSD. (b) Re-testing of social interaction behavior at T2-T4 of non-stressed and stressed animals (T2 p=0.0026, T3 p= 0.0045, T4 p=0.7145, 2-way repeated measures ANOVA and Bonferroni’s post hoc test). SI indices of control animals significantly decreased at time points T3 and T4 compared to T2 (T2 vs T3 p = 0.008, T2 vs. T4 p = 0.005, T3 vs. T4 p > 0.999, 2-way repeated measures ANOVA followed by Bonferroni’s posthoc test) (c) Group dynamics of social interaction behavior in stressed mice at three different time points (T2-T4). SI scores were significantly lower at T3 compared to T2 and recovered at T4 (T2 vs. T3 p = 0.017, T2 vs. T4 p = 0.977, T3 vs. T4 p = 0.197, 2-way repeated measures ANOVA followed by Bonferroni’s post-hoc test). (d) Longitudinal trajectories of stressed animals at T2-T4 of the 5 worst performers chosen at T2. (e) Longitudinal trajectories of stressed animals at T2-T4 of the 5 best performers chosen at T2. Data are shown as mean ± standard error of the mean and individual values.

Re-testing SI behavior in the same animal seemed to reduce the interest of the animals in interacting with the CD1 mouse in the cylinder in both groups at the later time points (stressed and non-stressed, T3, T4) as indicated by the significant main effect of time in a 2-way repeated measures ANOVA (F_2,70_= 9.42, p = 0.0002). This re-testing effect was avoided in the design of the first experiments (Fig. 1) when different groups of animals were tested at the different timepoints. Nevertheless, looking at group behavior and comparing non-stressed with stressed animals at T2 and T3 the stressed animals performed significantly less well in the SI testing than the control group (F_1,35_= 21.11, p <0.001, post-hoc comparisons are shown in Fig. 2b). With regard to the longitudinal changes of SI scores in stressed animals, we observed a significant drop in the performance at T3 compared to T2 (Fig. 2c). Interestingly, slight (not statistically significant) recovery in social behavior at time point T4 compared to T3 was observed (Fig. 2b, c) so that the difference of the stressed group compared to the behavior of the control animals was extinguished (Fig. 2b).

In order to take a closer look at individual behavior we selected the best and the worst 5 performers at T2 and displayed individual behavior tracks on separate graphs (Fig. 2d, e). Of the 5 worst performers, except for animal 6455 that reached an even smaller SI score at T4 than at T2, the other four animals improved SI behavior by T4 (Fig. 2d). Two of the five animals (6459, 6485) reached an SI score of above 100 at T4, three animals still stayed beneath 100 and therefore still showed (slightly) affected behavior. Animals with an SI score of above 100 have previously been defined as resilient performers (5). Three of the five animals (6455, 6485, 6488) reached an SI score peak at T3 followed by a drop of SI behavior afterwards (Fig. 2d).

Contrary to these results, of the five best performers at T2, three (6475, 6476, 6482) dropped in SI behavior at T4 beneath 100 and two (6473, 6483) kept their good score or even improved (6473) and reached scores above 100 and thereby met the criteria of resilience (Fig. 2e). All five animals dropped in SI behavior at T3 compared to the T2 scores followed by a further drop in three animals (6475, 6476, 6482) until T4 (Fig. 2e).

These data suggest that recovery in SI behavior is reached at T4 at the group level. However, individual behavior follows a dynamic trajectory throughout the analyzed time points.

### Longitudinal evolvement of cognitive behavior changes in individual animals

NOR testing with 24 hours between sample and test phase was carried out in the same group of animals directly after SI testing at T2-T4 (Fig. 2a). Confirming the previous analysis (Fig. 1f, g) this showed that cognitive behavior was lower at T2, improved 3 weeks after CSD (T3) and dropped again at T4, although changes were not statistically significant (Fig. 3a). These changes could be influenced by a significant drop in exploration behavior at T4 compared to T2 (supp. Fig. 5). However, the difference in DI scores between time-points was not significant even after adjusting for the effect of exploration time in an analysis of covariance.

**Figure 3.**
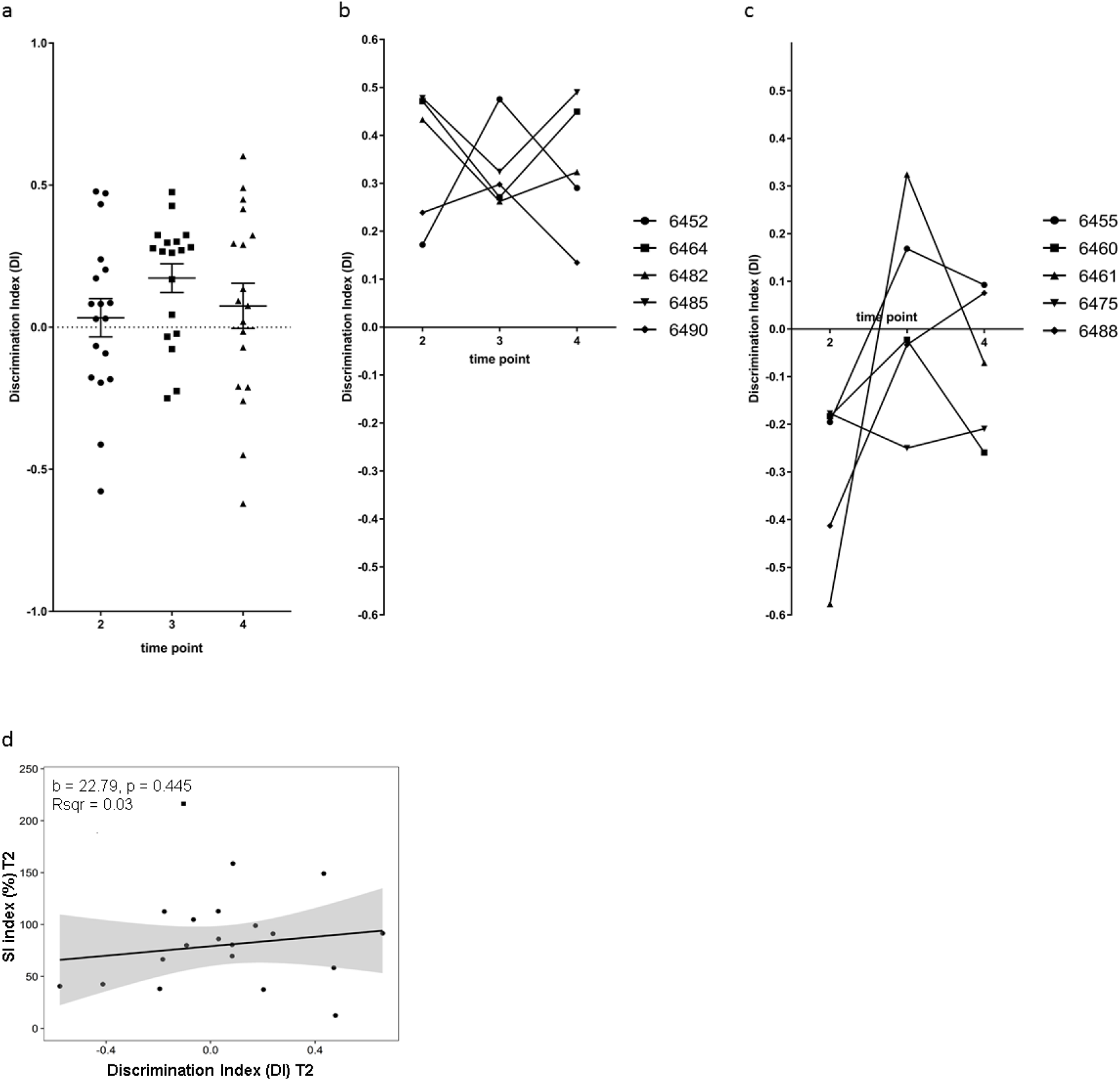
Longitudinal development of novel object recognition behavior after CSD stress. (a) Group dynamics of novel object recognition behavior in stressed mice at three different time points (T2-T4). DI scores were low at T2 and T4 and had temporarily improved at T3 after CSD exposure. (T2 vs. T3 p=0.4376, T2 vs. T4 p=>0.9999, T3 vs. T4 p=0.914, ANOVA and Bonferroni’s post hoc test) (b) Longitudinal trajectories of stressed animals at T2-T4 of the 5 best performers chosen at T2. (c) Longitudinal trajectories of stressed animals at T2-T4 of the 5 worst performers chosen at T2. (d). Linear regression analysis using DI scores of stressed animals at T2 as independent and SI scores as dependent variable revealed no significant results for the regression coefficient b and the amount of variance explained as indicated by R-squared. The gray area around the regression line corresponds to the 95% confidence band. One animal was excluded from the regression analysis due to being a statistical outlier based on its SI score (shown as a square on the scatter plot). Non-linear trends in the relationship were investigated using a general additive model. Goodness of fit was not improved as confirmed by a likelihood ratio test, therefore a linear model was fitted to the data. Data are shown as mean ± standard error of the mean and individual values.

Again, we looked at individual behavior and picked the best and worst 5 performers at T2 and tracked them individually (Fig. 3b, c). For all animals (best or worst performers) novel object recognition changed between T2 and T3. While 2 of the best performers (6452, 6490) performed better at T3 and then dropped again at T4, the three others dropped in performance at T3 and then improved again at T4 (Fig. 3b). Three mice performed less well at T4 compared with T2 (6452, 6482, 6490). All animals of the best performer group at time point T2 still performed above average at T4 past CSD (Fig. 3b).

Four of the five animals in the group of the worst performers at T2 improved considerably at T3 (Fig. 3c). While three of these (animals 6455, 6460, 6461) dropped again in their performance at T4, four still improved in object recognition at T4 compared to T2. Only one animal of the worst performer group kept the low performance level (6475) from T2 (Fig. 3c).

Finally, we investigated if novel recognition performance can predict social interaction behavior at T2 using a linear regression analysis (Fig. 3d). Results did not reveal a significant association between DI and SI scores, which confirmed results from the previous experimental setup (Fig. 1j).

### Correlation between behavior at T2 and T4

Last we asked if behavior of animals at T4 can be predicted from behavior shown at T2. To answer this question, we performed linear regression analysis using DI and SI scores at timepoint T2 as independent variables and scores at T4 as dependent variables. SI scores at T2 did not predict SI scores at T4 (Fig. 4a). Furthermore, DI scores at T2 were not significant predictors for either DI or SI scores at timepoint T4 (Fig. 4b, c).

**Figure 4.**
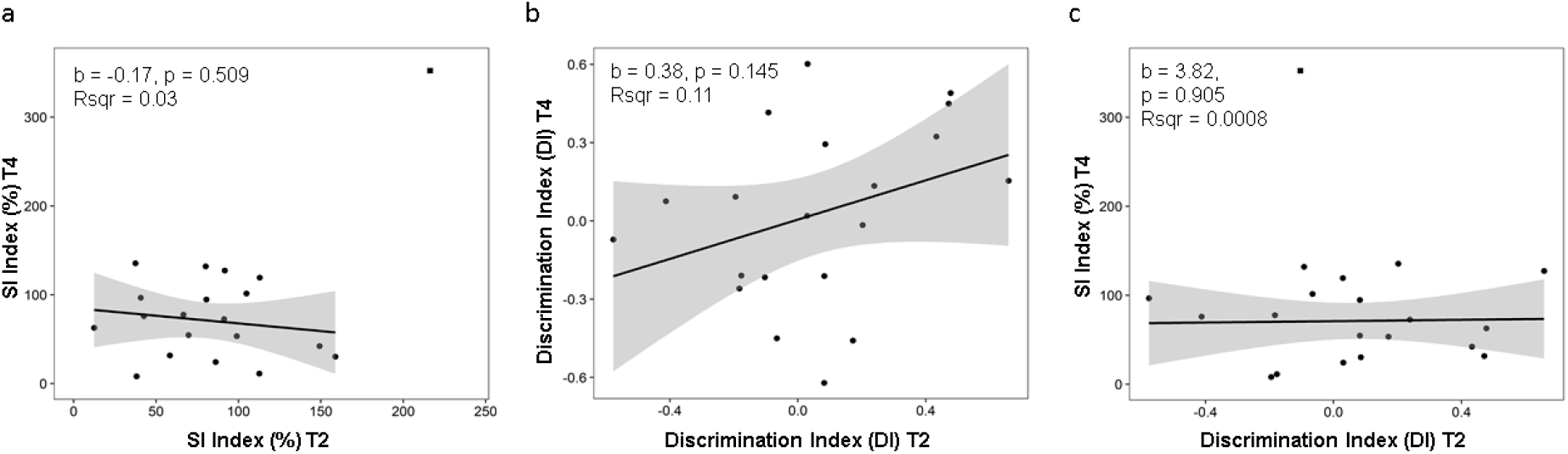
Linear regression analysis for social interaction and novel object recognition. (a) SI scores at timepoint T2 did not significantly predict SI behavior at T4. (b) DI scores at T2 did not significantly predict DI scores at T4. (c) DI scores at T2 did not significantly predict SI scores at T4. One animal was excluded from the regression analysis of SI scores due to being a statistical outlier (shown as a rectangle on the plots). For each regression analysis, the regression coefficient b and the amount of variance explained (R-squared) are shown. The gray area around the regression line corresponds to the 95% confidence band. Non-linear trends in the relationship were investigated using a general additive model. Goodness of fit was not improved as confirmed by a likelihood ratio test, therefore a linear model was fitted to the data.

These data suggest that behavior trajectories of individual animals are highly dynamic in response to chronic social defeat stress. Predictions of long-term behavior at an early time point of 7 days post exposure are not possible.

## Discussion

We show here that, in addition to SI behavior, also novel object recognition in a test setting with 24 hours between sample and test phase is impaired in C57BL/6J mice 7 days after the last defeat of a CSD exposure. We could not find a correlation between impaired SI and novel object recognition behavior at the 7 days (T2) time point. This argues for an independent influence of CSD on social behavior and object memory. This is also supported by the different longitudinal courses both behavior traits took after CSD exposure. Novel object recognition impairment was only seen at T2 but not at T1 and then seemed to go through a period of improvement at T3 but showed impairment again at T4. SI behavior by contrary seemed to be affected already at T1 and remained impaired until T3. At T4 however recovery of SI behavior seemed to occur when looking at group dynamics.

Several brain regions have been assigned to be involved in social avoidance and depressive behavior after CSD exposure including the prefrontal cortex, amygdala, the ventral tegmental area, the ventral hippocampus and the nucleus accumbens (14–18). Novel object recognition with 24 hours between test and sample phase has been assigned to the hippocampus (19, 20). Our data suggest that stress influences both circuitries independently. Furthermore, it shows that memory aberrations after stress do not influence social interaction behavior and that connectivity between these two behavior domains is low.

Group performance after CSD in the social interaction test started being affected already at T1, stayed low at T2 and T3 but increased again at T4. Interestingly, social interaction activity was reduced by both, stressed and non-stressed animals after the first testing. This effect might be due to a decrease in interest of no longer naïve animals. Still, significant differences in group performance were visible between control and stressed animals at T2 and T3 (Fig. 2b), while at T4 control and stressed groups performed similarly. Our data argue for underlying molecular and functional effects that establish during or quickly after CSD exposure and that cause an adaptation process finally leading to full recovery at T4. However, individual tracks draw a much more complicated and dynamic picture.

Looking at the best performing animals at T2 after stress exposure three of five dropped significantly at T4, far beneath the 100 SI-score level, which is thought to be the threshold for resilient versus susceptible behavior. Four out of five from the group of the worst performers on the other hand significantly recovered between T2 and T4 and two reached values above 100 SI scores, thereby meeting criteria of resilience. These observations point towards a dynamic process. Instead of defining only two groups, resilient and susceptible at a time point relatively close to CSD exposure, our data give evidence for the existence of four groups: (i) resilient shortly after CSD and still resilient at T4, (ii) resilient shortly after CSD and developing impairments later, (iii) susceptible shortly after CSD and, while improving, still impaired in SI behavior at T4 and (iv) susceptible shortly after CSD but having recovered into resilience at T4.

Novel object recognition is unaffected at T1, declines at T2, then goes through a period of recovery at T3 and finally tends to go down again at T4 after CSD exposure. This suggests that molecular mechanisms influencing NORT remain stable for a couple of days after exposure before they drop. This might hint at the involvement of proteins or mRNA with a long half-life time requiring degradation before an effect can be recognized. At the same time impairment in novel object recognition undergoes temporary changes leading to recovery at T3. This suggests short-term homeostatic processes kicking in at the time period between T2 and T4 trying to compensate for effects caused by chronic social defeat stress. Alternative to repair mechanisms of molecular changes caused by CSD stress, the recovery could also be due to active adaptation processes that are able to e.g. compensate quickly for changes in gene / protein expression. The drop in object recognition behavior at T4, however, argues for a highly dynamic and complex homeostatic process that has varying effects on behavior with time.

Interestingly, regression analysis revealed that animal behavior 24 hours or 7 days after CSD is not predictive for behavior after 42 days in both behavior domains, social interaction and novel object recognition (Fig. 4). In humans, PTSD (post traumatic stress disorder) is a pathological reaction to stress-trauma that has a substantial effect on life quality of affected patients. PTSD is defined as a combination of symptoms from four symptom clusters that develop after a traumatic event and hold for longer than a month (21, 22). Comparable to what we see in mice, human subjects can show an immediate or quick reaction to stress as an acute stress disorder (less than 48 hours after the event) or an adjustment disorder (more than a week and less than a months after the event) that might include psychological and cognitive symptoms (22). They then either recover or continue showing and developing symptoms possibly resulting in PTSD. Although life spans in humans and mice differ significantly, molecular mechanisms that are likely to underlie stress reactions including protein and mRNA stability, transcription and translation and epigenetic marking follow similar time scales. A mouse model for stress induced resilience or susceptibility should consider this and look longitudinally at long-term behavior. The fact that our data do not show correlations between short- and long term behavior after CSD, suggests that differentiation between resilient and susceptible outcome should not be made shortly after stress exposure in the CSD model used here. We argue that only by including longitudinal observations behavior reactions of adult animals exposed to chronic social defeat stress are a plausible model for the complex impairment of mental function occurring in human subjects after trauma. It will be interesting to see how behavior develops in adult animals longitudinally in other models of chronic stress.

## Acknowledgement

The work was funded by the DFG (Deutsche Forschungsgemeinschaft), CRC 1193.

---

**Supplementary Figure 1:**
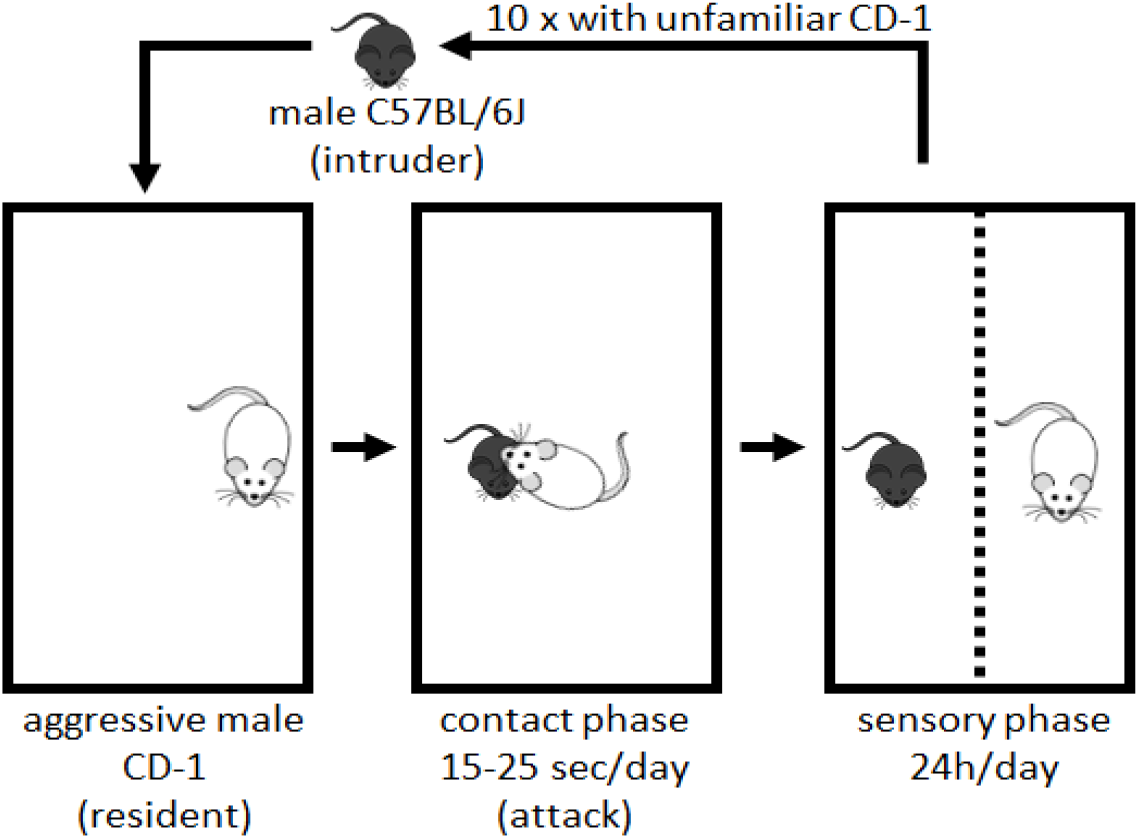
Schematic representation of the CSD.

**Supplementary Figure 2:**
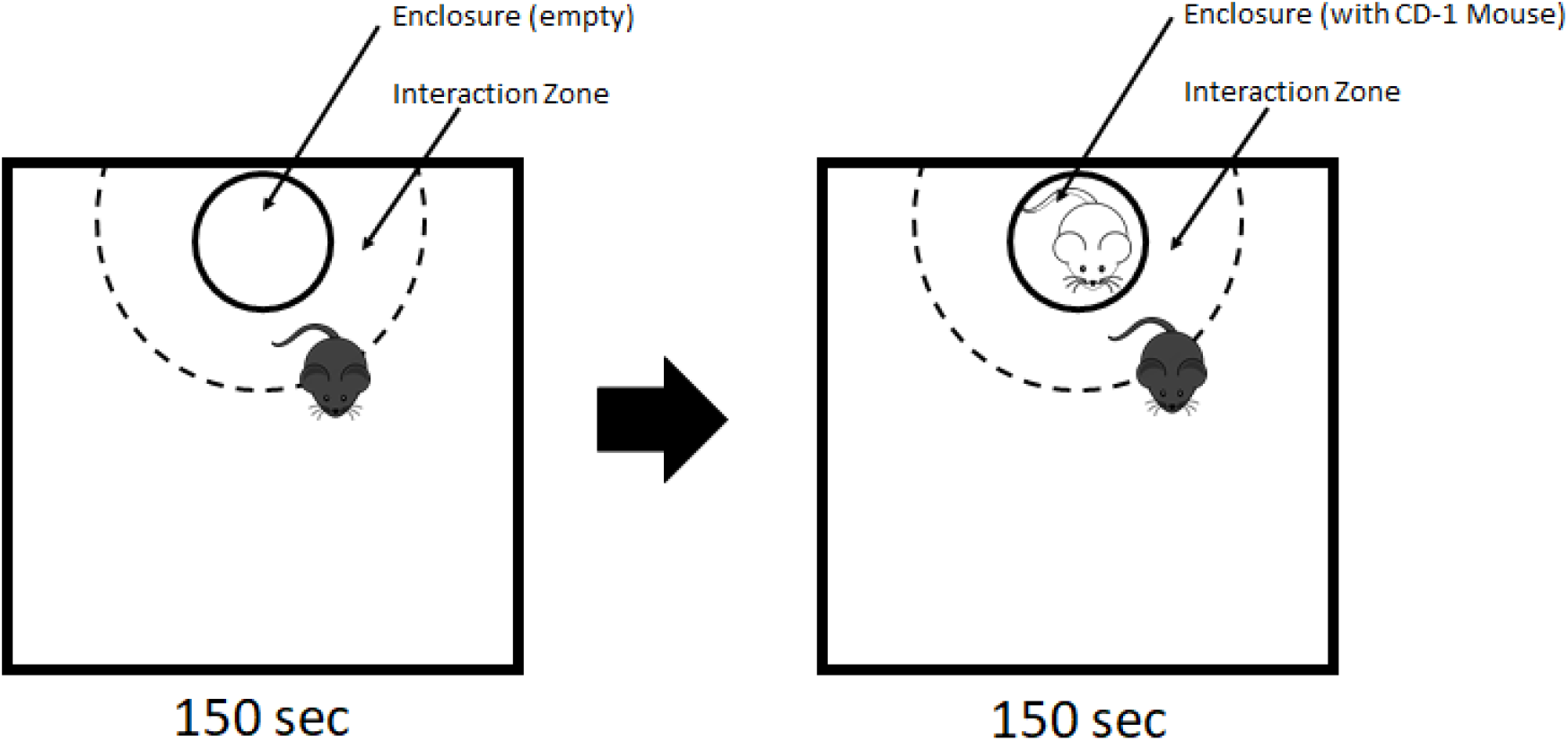
Schematic representation of the SI test.

**Supplementary Figure 3:**
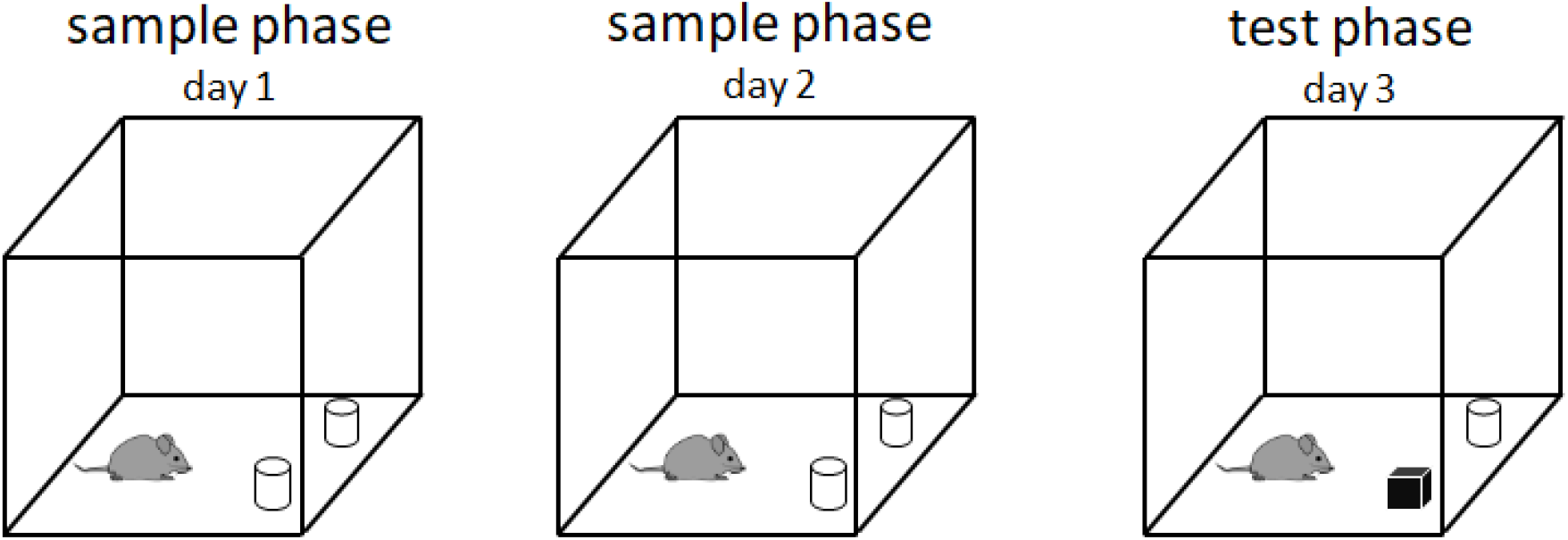
Schematic representation of the NOR test.

**Supplementary Figure 4a-c:**
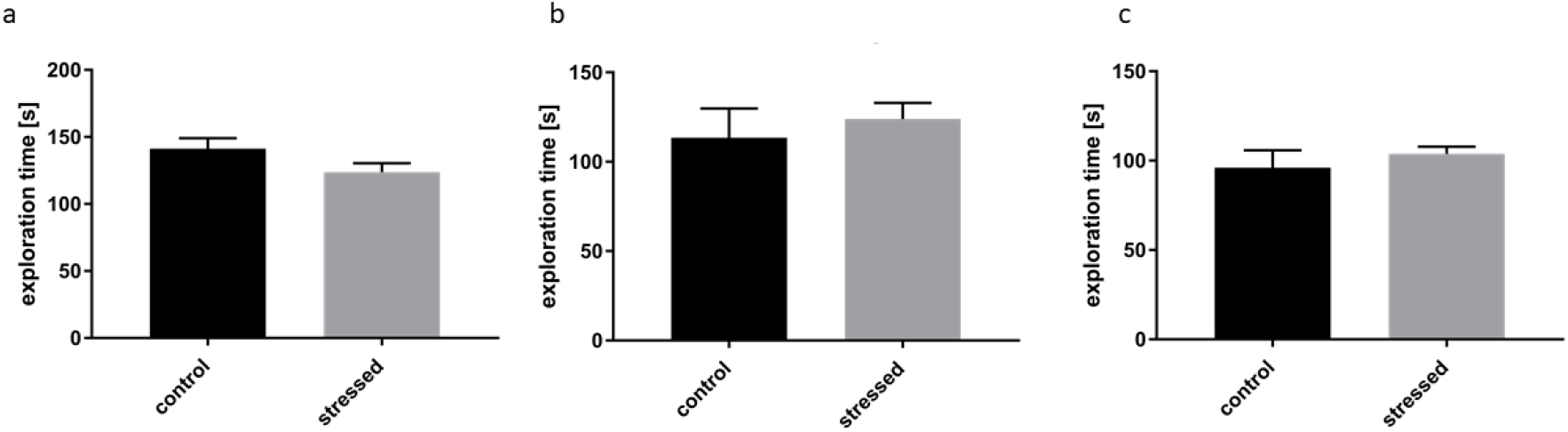
Exploration times measured in novel object recognition testing of animals according to Figure 1e-g. (a) 24 hours post exposure. (p=0.1641, unpaired t-test with Welch’s correction) (b) 8-10 days post exposure. (p=0.5838, unpaired t-test with Welch’s correction) (e) 22-24 days post exposure (p=0.7836, unpaired t-test with Welch’s correction). Data are shown as mean + standard error of the mean.

**Supplementary Figure 5:**
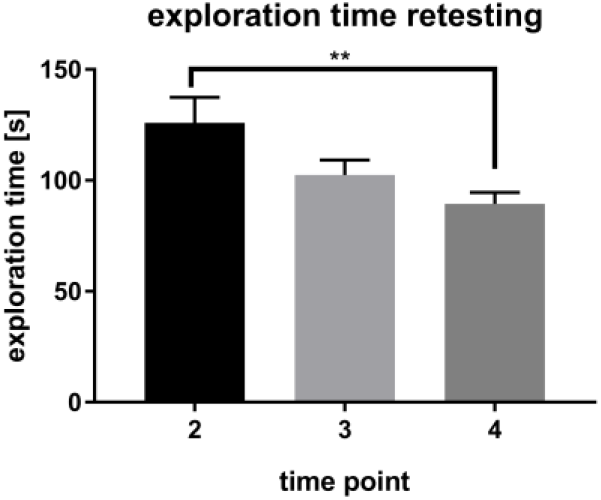
Exploration times measured in novel object recognition testing of animals in figure 3 at three different time point. T2: 8-10 days post exposure. T3: 22-24 days post exposure. T4: 43-45 days post exposure (T2 vs. T3 p=0.1504, T2 vs. T4 p=0.0089, T3 vs. T4 p=0.8118, ANOVA and Bonferroni’s multiple comparisons test). Data are shown as mean + standard error of the mean.

